# Host Microbiome Richness Predicts Resistance to Disturbance by Pathogenic Infection in a Vertebrate Host

**DOI:** 10.1101/158428

**Authors:** Xavier A. Harrison, Stephen J. Price, Kevin Hopkins, William T. M. Leung, Chris Sergeant, Trenton W. J. Garner

## Abstract

Environmental heterogeneity is known to modulate the interactions between pathogens and hosts. However, the impact of environmental heterogeneity on the structure of host-associated microbial communities, and how these communities respond to pathogenic exposure remain poorly understood. Here we use an experimental framework to probe the links between environmental heterogeneity, skin microbiome structure and infection by the emerging pathogen *Ranavirus* in a vertebrate host, the European common frog (*Rana temporaria*). We provide evidence that environmental complexity directly influences the diversity and structure of the host skin microbiome, and that more diverse microbiomes are more resistant to perturbation associated with exposure to *Ranavirus*. Our data also indicate that host microbiome diversity covaries with survival following exposure to *Ranavirus*. Our study highlights the importance of extrinsic factors in driving host-pathogen dynamics in vertebrate hosts, and suggests that environment-mediated variation in the structure of the host microbiome may covary with observed differences in host susceptibility to disease in the wild.

## Introduction

Animals host to diverse communities of microbes, collectively referred to as the microbiome. Host-associated microbes are increasingly recognized as a crucial component of the immune response of multicellular organisms (Ford and King 2016), and evidence from a diverse suite of host species has identified these symbiotic microbes as key mediators infection by pathogens (Spor et al 2011; Koch & Schmid-Hempel 2011; Ford & King 2016; King et al 2016). The microbiome can interact with pathogens either indirectly by modulating the host immune response (Rollins-Smith 2009), or directly by releasing anti-microbial compounds that kill the pathogen (King et al 2016). Crucially, though differences in host microbiome may predict variation among individuals in resistance to infection (Kueneman et al 2016), successful invasion by a pathogen may also disrupt the existing microbiome (Jani & Briggs 2014; Longo & Zamudio 2017). Understanding the factors that determine the strength of disruption of the host microbiome by pathogens is important; it represents a fundamental component of our knowledge of how pathogens and microbes interact, and is critical for predicting host responses to infection. For example, disruption of the host microbiome by potential pathogens may compromise important physiological functions, including immunity, that could increase the risk of invasion by additional pathogens or alter the dynamics of the primary infection.

The amphibian skin microbiome is rapidly becoming established as a model system for understanding the tripartite relationships between host, bacterial microbiome and pathogens (Jani & Briggs 2014; Longo et al 2015). Production of metabolites by skin-associated bacteria is a crucial component of immune defense against the lethal fungal pathogen *Batrachochytrium dendrobatidis* (*Bd*; Brucker et al 2008a,b; Harris et al 2009; Van Rooij et al 2015; Kueneman et al 2016). Fascinatingly, the production of anti-fungal metabolites by bacteria increases dramatically when they are co-cultured (Loudon et al 2014a), suggesting that microbiome-mediated host protection is likely a function of synergistic interactions among community members. Greater microbiome diversity may therefore offer increased protection from pathogens and diseases they cause (Kueneman et al 2016), but a major outstanding question is whether the susceptibility of the host-associated microbiome to perturbation by pathogens is also a function of its diversity. Though several studies have sought to measure the influence of pathogenic infection on host microbiome structure (Jani & Briggs 2014; Longo et al 2015; Longo & Zamudio 2017), all have assumed that the magnitude of disruption is uniform for all infected hosts and not modulated by their initial state. This assumption is poorly supported; there is strong evidence that host-associated microbial communities are heavily influenced by abiotic conditions (Caporaso et al 2011; Hyde et al 2016; Rebollar et al 2016; Jiminez & Sommer 2017), including the microbial diversity of the surrounding habitat (Costello et al 2012; Walke et al 2014). Thus, spatial or temporal variation in microbiome diversity mediated by the environment may modify the strength of interactions between the host microbiome and invading pathogens. There is now growing interest in the idea that environmental fluctuations may alter the susceptibility of hosts to pathogenic infection by driving changes in the microbiome (Chang et al 2016; Longo & Zamudio 2017). The diversity-stability hypothesis predicts that more diverse communities should be more resistant to disturbance, and several empirical studies support this hypothesis in plant community assemblages (McCann 2000; Costello et al 2012), but it is unclear whether this ecological theory is also relevant at the scale of host-associated microbial communities (Costello et al 2012). Furthermore, all studies to date have used the fungal pathogen *Bd* as an infectious agent, and comparative data on pathogen-mediated disruption of the microbiome from other pathogen groups are required to validate this theory.

Here, we use an experiment to test the influence of the complexity of the environmental bacterial reservoir on host microbiome diversity, and test whether the magnitude of disruption of the host microbiome by exposure to a pathogen is dependent on its diversity. In addition, we measure the association between host microbiome diversity and survival following exposure to a viral pathogen of the genus *Ranavirus* (family *Iridoviridae*) in European common frogs (*Rana temporaria*). Emerging infectious diseases (EIDs) caused by pathogens with broad host ranges like ranaviruses represent a significant threat to animal health, and are associated with global declines in biodiversity and population extirpations (Lips et al 2006; Price et al 2014; Rosa et al 2017). *Ranavirus* was responsible for multi-species amphibian declines in continental Europe (Price et al 2014) and for declines of the common frog in the United Kingdom (Teacher et al 2010).

We predicted that i) host microbiome diversity will increase in tandem with the bacterial community diversity of the environment; ii) that more diverse microbiomes will be more resistant to disruption by pathogenic infection; and iii) that host microbiome diversity will correlate with survivorship following exposure to an FV3-like ranavirus. To manipulate host microbiome complexity, we assembled experimental units that either contained a natural bacterial reservoir (*complex* habitats, containing a soil substrate and leaf litter) or lacked one (*simple* habitats, containing sterile stony terrestrial substrates and no leaf litter). Both groups comprised 8 units containing 6 individuals each (n= 48 individuals per habitat treatment). After 14 days in experimental habitats, we exposed half the individuals from each group to either *Ranavirus* or a sham control to measure i) habitat-dependent effects of disruption to the host microbiome by a pathogen and ii) habitat-dependent mortality following exposure to *Ranavirus* or the negative control.

## Methods

### Ethical statement

All experimental procedures and husbandry methods were approved by the ZSL Ethics Committee before any work was undertaken and was done under licensing by the UK Home Office (PPL 70/7830). Animal health and welfare was monitored daily during both the rearing and experimental periods and all animals were fed *ad libitum* (Tetra Tabimin for tadpoles, small crickets for metamorphosed frogs) throughout.

### Experimental Protocol

Animal Rearing: *R. temporaria* metamorphs were reared from tadpoles hatched from clutches sourced from UK garden ponds. Animals that completed metamorphosis were cohoused in large groups (no more than 40 per enclosure) in 460 × 300 × 170mm Exo Terra Faunaria containing cleaned pea gravel, a large cover object and sloped to accommodate a small aquatic area. Experimental animals were haphazardly selected from four group enclosures.

Preparation of habitat treatment enclosures: The general layout of both habitat types was shared in that they both contained a filled, plastic PCR tip box forming a terrestrial platform elevated above an aquatic area filled with aged tapwater, a cover object located on the terrestrial platform, and autoclaved pea gravel formed into a slope leading from the aquatic area to the platform, with each replicate housing six recently metamorphosed frogs. The two key differences were that i) the terrestrial platforms in complex habitats contained organic compost as a substrate, whilst the terrestrial platforms in simple habitats contained standard and autoclaved pea shingle and ii) leaf litter collected from Regents Park, London, was added to the aquatic area in the complex habitats. complex habitat enclosures were left uncovered and outdoors for two weeks prior to the start of the experiment, while simple habitat enclosures were prepared the day before frogs were transferred into replicates. During the experiment, uneaten cricket corpses were removed from simple habitat enclosures, but left in complex habitat enclosures.

Experimental procedure: Following rearing in an outdoor facility, animals were moved to a procedure room and housed individually for seven days in Perspex boxes with a cover object and damp paper towel as substrate to acclimatize. We randomly allocated individuals to complex or simple treatments for 14 days before sham or ranavirus exposures. Samples for 16S metagenomics were collected from living animals immediately preceding transfer to experimental units and at the end of the 14 day period by swabbing the skin of the body and limbs of frogs. Environmental swab samples (two per experimental unit, one terrestrial and one aquatic) were also collected on day 14 preceding pathogen exposure procedures. Terrestrial swabs were taken by running the swab over the terrestrial substrate and inside the cover objects twice. Aquatic swabs were taken by submerging the swab in the aquatic portion of the tank

Microcosms were randomly assigned to disease treatment group (ranavirus or sham) using a script written in R. Previous to this, *Ranavirus* (FV3-like isolate RUK13, Price et al. 2016) was cultured in EPC cells at 27°C, harvested after cell layer had completely cleared, subjected to three rounds of freeze-thaw and then cleared of cells and cellular debris by centrifugation at 800g for ten minutes and discarding the cell pellet. Virus titre was estimated using a 50% Tissue culture Infective Dose assay (TCID_50_) and calculated following the method of Reed and Muench (1938). Sham exposure media was produced by harvesting a pure culture of EPCs and harvesting the supernatant after the same 800g, ten minute spin. For exposures, animals were transferred as experimental units groups to 90 mm petri dishes containing 19 mL of aged tap water. Depending on treatment, either 1 mL of stock virus culture at 2 × 10^6^ TCID_50_/mL (giving a final exposure concentration of 1 × 10^5^ TCID_50_/mL) or 1 mL of sham media was added to the petri dish. Animals were exposed in petri dishes for six hours before being returned to their habitat treatment enclosures.

Frogs and experimental unit environments were swab sampled again for 16S metagenomics on day 2 post-exposure. We used daily health and welfare checks throughout the experiment to monitor survival rates. We also used daily checks to monitor for signs of disease commonly associated with ranavirosis (see below). We ended the experiment on day 30 when all frogs appeared physically healthy and when mortality had subsided.

### 16S Metagenomic Sequencing

16S metagenomic library preparation was carried out using a modified version of the protocol detailed in Kozich *et al* (2013) that amplifies the v4 section of the 16S rRNA gene. Sequencing was performed using 250bp paired-end reads on an Illumina Miseq using a v2 chemistry 500 cycle cartridge. Raw 16S metagenomic sequence data were analysed using *mothur* v1.36.1(Schloss et al 2009) and exported as a ‘biom’ object to be read directly into the R package *phyloseq* (McMurdie & Holmes 2013). A detailed workflow is provided in supplementary methods.

### Statistical Analyses

We calculated estimates of microbial community richness and structure after rarefying to even sequencing depth to remove bias caused by differences in ‘sampling effort’ across libraries of different sizes (Rarefied Sequence Analysis). We also analysed differences among treatment groups using the overdispersion-corrected analytical framework on unrarefied sequence data to use all sequencing reads (Differential Abundance Analysis; McMurdie & Holmes 2014). All statistical analyses of microbiome data were performed on 85 individuals (42 in Simple Habitats and 43 in Complex habitats). Eight individuals were censored because they died before the pre-infection Day 14 microbiome swab. Two swabs were excluded because of a labeling error meaning they could not be unambiguously attributed to a particular block, and 1 sample was excluded because it fell below the 10,000 read threshold for library rarefaction (n = 6447 reads). This latter sample was not excluded from the ‘Differential Abundance’ analysis (see below). A fully reproducible workflow for all analyses is provided as an R markdown document.

<u>Rarefied Sequence Analyses:</u> We followed the protocol in Longo & Zamudio (2017) and filtered out all OTUs with <100 reads (n = 7175), leaving 1053 OTUs used in downstream analysis. Results using all OTUs were similar and not shown here. All libraries were rarefied to 10,000 reads per sample. We used the Chao1 metric to compare differences in richness among treatments, and the R package *vegan* (Oksanen et al 2015) to visualize microbial community structure differences across samples using NMDS ordination. NDMS ordinations were performed with k = 2 and yielded stress values <0.13. We used *adonis* to test for significant differences in community structure among groups using permutational ANOVA. To test the effects of habitat complexity on microbial community structure after 14 days (pre-infection dataset), we fitted habitat treatment as a single predictor; and ii) to test the effect of both ranavirus infection and habitat complexity on community structure, we fitted both infection treatment, habitat treatment and their interaction as predictors.

<u>Differential Abundance Analyses:</u> We fitted similar models to the *adonis* models above in the R package DESeq2 (Love et al 2014). DESeq2 models provide quantitative estimates of differences in bacterial abundance between experimental treatments whilst simultaneously controlling for overdispersion introduced by differences in library size using a Negative Binomial mixture model (McMurdie & Holmes 2014). This allows the identification of significantly differentially abundant OTUs between treatments whilst avoiding the bias introduced by rarefying libraries to even size. Significant OTU abundances between treatments were quantified using Wald tests with p values corrected for multiple testing.

<u>Permutational Test of Differential Abundance:</u> To test if differences in susceptibility to disturbance recovered by DESeq2 analysis were likely to have arisen by chance, we conducted a permutation test on the post-exposure data. For each iteration we randomly assigned all individuals in a block to a habitat complexity (simple/complex) and exposure (ranavirus / sham) treatment, and reran the DESeq model ‘ ∼ habitat*disease’. At each iteration, we stored the number of differentially expressed OTUs between ranavirus and sham exposures within each habitat type (i.e. comparing ranavirus-exposed to sham-exposed individuals in complex habitats). We ran a total of 1000 permutations to derive a null distribution for the number of differentially-abundant OTUs for each habitat type.

### Survival Data

To investigate the effects of habitat complexity and ranavirus infection on survival, we fitted mixed effects Cox models using the ‘coxme’ package (Theriot et al 2014) in R v3.2.4 (R Core Team 2017). All models contained a right-censored survival object as a response, comprising a two-column vector of ‘time of event’ and an indicator variable representing whether mortality was observed for that individual (1) or not (0). Block ID was fitted as a random effect to control for block effects. The most complex model contained habitat treatment, infection treatment, and their interaction as fixed effects. We fitted all nested models and ranked them by AICc. Models that were more complex versions of a model with better AIC support were removed following the nesting rule (Richards 2008). We used survival data from Day 15 (day of exposure) to examine the effect of ranavirus on survival as there was a degree of background mortality prior to Day 15 (n=8 of the 96 individuals), giving a total sample size of 88 individuals.

### Signs of Disease Data

We used the R package *MCMCglmm* (Hadfield 2010) to analyse visible signs of infection for the individuals in the RV+ treatment group. Total sample size for this analysis was 42 individuals alive at the time of exposure to ranavirus. We analysed both i) probability of observing signs of disease and ii) severity of disease signs, where visible signs of disease typical of ranavirosis were scored on a 3 point scale (0: no visible signs; 1: visible redness on limbs and venter, including subcutaneous petechial hemorrhages, and body oedema; and 2: signs of ulceration on limbs, body, digit tips). For i) we fitted a Binomial error structure where signs of disease was coded as a 0/1 binary variable. For ii) we fitted an ordinal model with the categorised disease status as a response. Both modes included experimental units ID as a random effect. For both models chains were run for a total of 500,000 iterations following a burnin of 100,000 iterations with a thinning interval of 100. We assessed convergence using the Geweke diagnostic (Geweke 1992) (all z scores > - 0.4 & <0.8). We used uninformative priors for the random effects of box, and fixed the residual variance to 1 in both cases as neither Bernoulli nor ordinal models can estimate the residual variance.

## Results

### Habitat complexity predicts host skin microbiome community structure

Frogs inhabiting complex habitats had significantly greater skin bacterial alpha diversity than those reared in simple habitats (complex Chao1 index = 419.22 vs simple = 189.3; p_RAND_ < 0.001, Fig. 1a). This effect occurred despite all frogs having similar bacterial richness at the start of the experiment when housed individually (Online Supplementary Figure S1). Frogs in complex habitats possessed 682 unique Operational Taxonomic Units (OTUs) not detected in either the terrestrial or aquatic substrates, compared to 283 for frogs in simple habitats (Supplementary Figure S2). Frog skin bacterial richness increased in tandem with the bacterial richness of the environment (Pearson’s correlation t_14_ = 6.67, p<0.001, Supplementary Figure S3). Non-metric multidimensional scaling (NMDS) of individual frog skin microbial communities supported this pattern, with clear separation based on habitat type (Fig. 2a). Community composition was significantly different across habitat types when controlling for block effects (PERMANOVA simple vs complex p<0.001 r^2^ = 13.8%). Differential Abundance Analyses using DESeq2 on the unrarefied dataset identified 383 OTUs that were significantly different in abundance between complex and simple habitats after correction for multiple testing. Of these, 310 were significantly more abundant in complex habitats, whereas 73 were significantly more abundant in simple habitats (Fig. 3). In both simple and complex habitats, individual frogs possessed skin microbial community structures (beta diversity) that were distinct from the bacterial signatures of the terrestrial and aquatic areas in the experimental units (Supplementary Figure S4).

**Figure 1.**
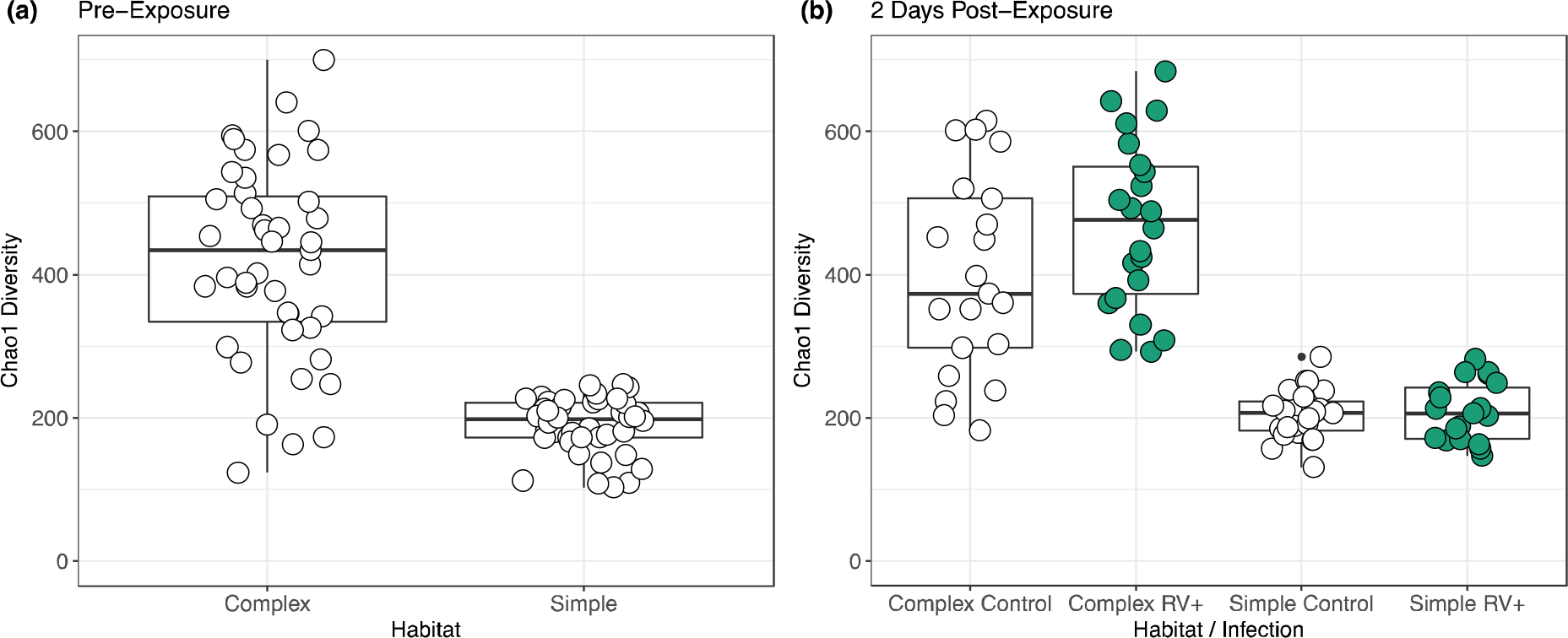
The effects of habitat complexity on common frog skin microbial community alpha diversity using the Chao1 richness estimator. (a) Chao1 diversity of 85 common frogs split across two habitat treatments differing in the richness of the microbial reservoir in the environment (complex and simple; n=43 and 42 per treatment, respectively). Frogs were kept in these treatment conditions for 14 days before quantifying microbial communities. (b) Frogs were exposed to either ranavirus (RV+) or a sham control (Control) and microbial communities were assessed 48 hours post-exposure. Raw data are displayed as points and jittered for display purposes.

**Figure 2.**
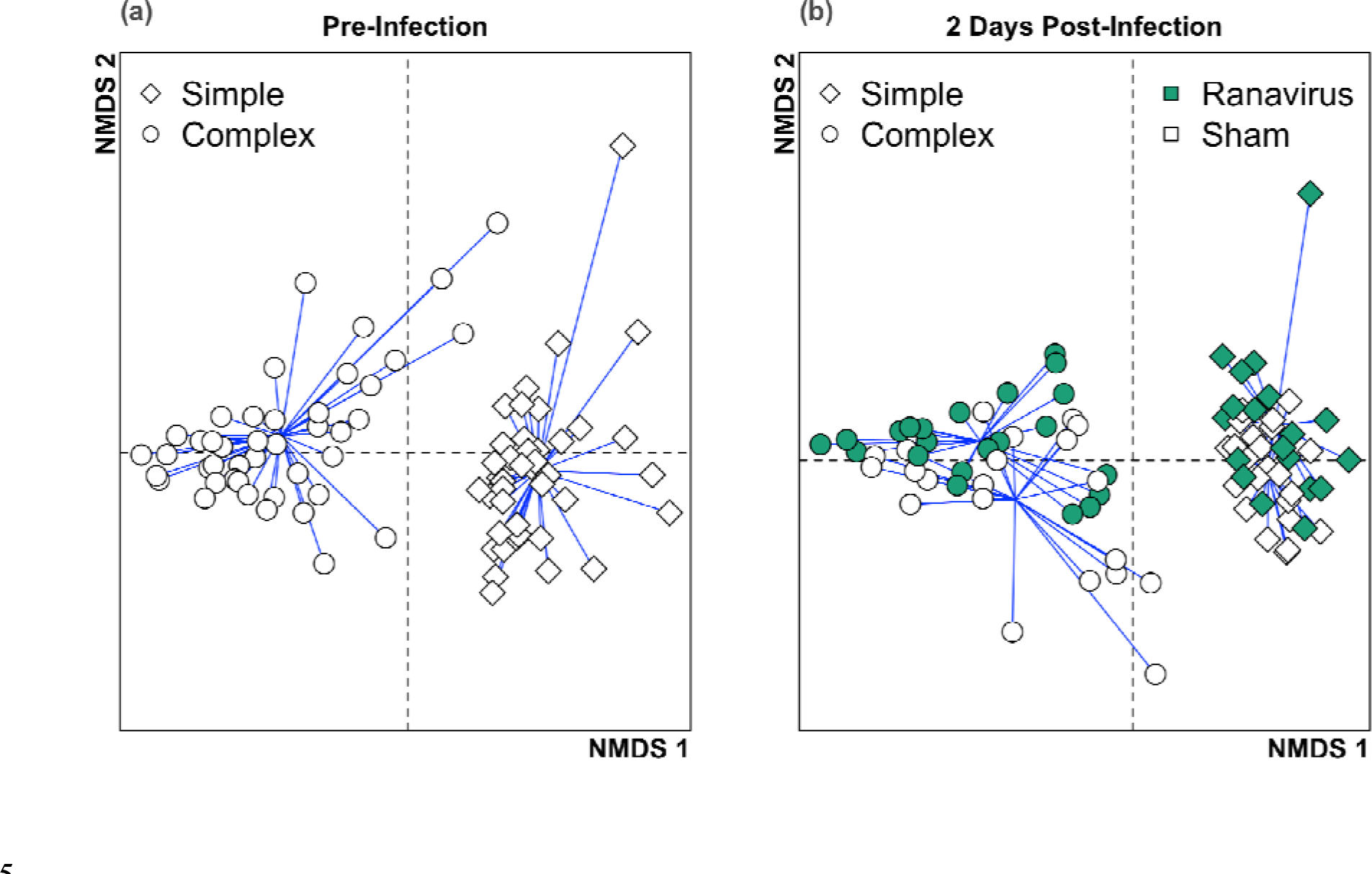
The effect of habitat complexity and exposure to Ranavirus on common frog skin microbial community structure. (a) Beta diversity differences among common frogs living in habitats with either high (complex) or low (simple) environmental bacterial richness. Adonis analysis revealed was a significant difference in the community structures of the two habitat treatments. (b) Half of each habitat were exposed to either ranavirus (green points) or a sham control (white points). Data presented in (b) were measured 48 hours post-exposure. Adonis analysis revealed that exposure to ranavirus perturbed the microbial community structure of both habitat type.

**Figure 3.**
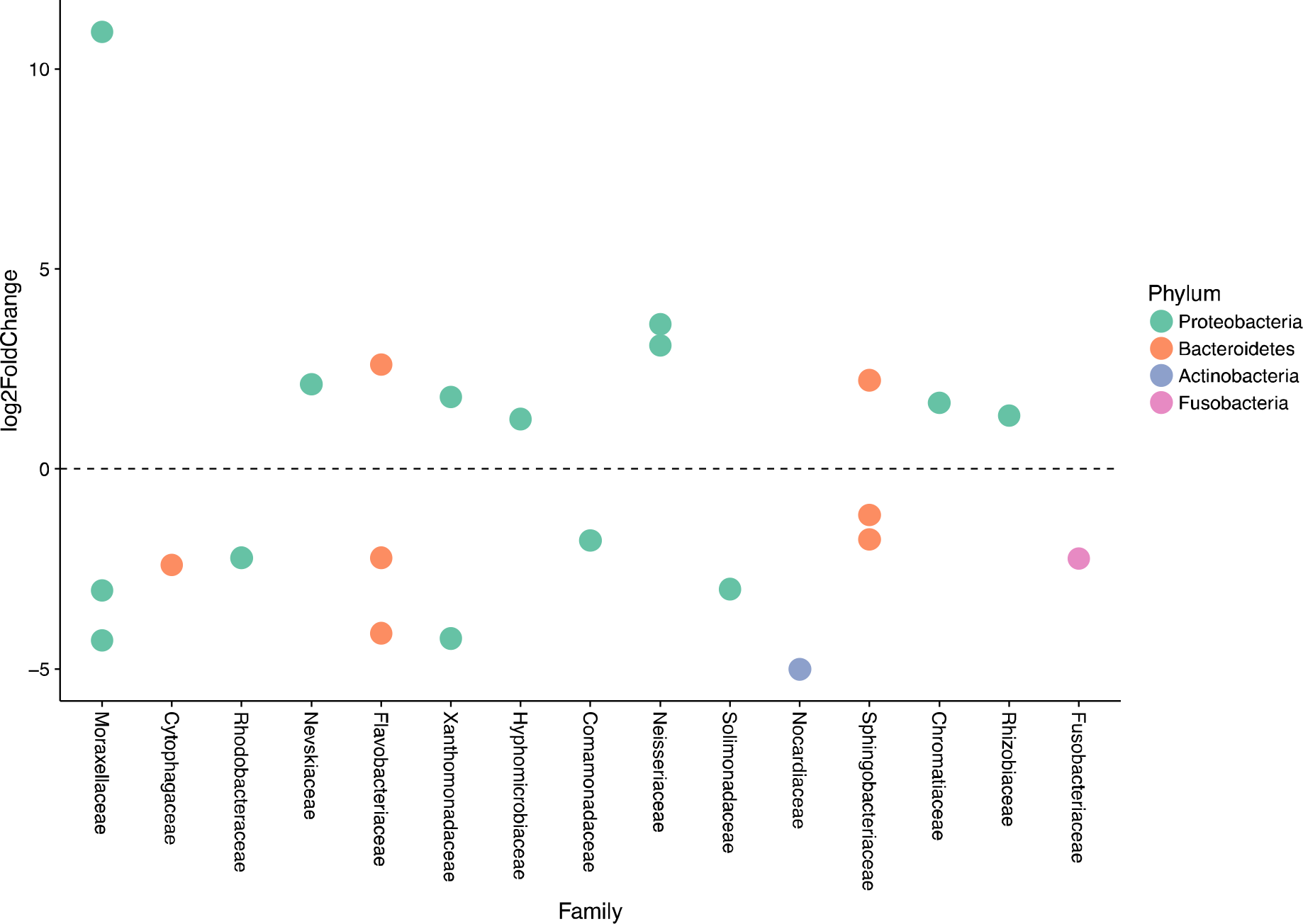
Differential abundance of 23 OTUs from the top 50 most abundant OTUs in common frog skin microbiome after 14 days in habitat treatments. Differences are for frogs in complex habitats as compared to frogs in simple habitats (positive log2FoldChange OTUs are enriched in complex habitats).

### Infection with *Ranavirus* Elicits Habitat-Dependent Disruption of the Host Microbiome

48 hours post exposure, there was no discernible effect of ranavirus exposure on the alpha diversity of frogs (p_RAND_ = 0.6), but the significant effect between habitats remained (p_RAND_ < 0.001, Fig. 1b). NMDS ordination at the individual level revealed subtle shifts in the centroids of *Ranavirus*-exposed frogs within habitat relative to the negative controls (Fig. 2b). Community composition of ranavirus-exposed frogs was significantly different from the controls whilst controlling for block ID (PERMANOVA, habitat p<0.001 r^2^ = 13.79%; exposure p<0.001 r^2^=3.5%). Differential Abundance Analyses supported these patterns and revealed more subtle effects, where exposure to *Ranavirus* caused shifts in microbiome community structure dependent on habitat treatment (Fig. 4). Frogs in complex habitats exhibited relatively low levels of change, with equal levels of increase and decrease in OTU abundance of 3 phyla relative to controls. Conversely, exposure to*Ranavirus* in simple habitats resulted in broader changes in abundance of bacterial community members (31 significantly different OTUs in simple habitats compared to 11 in complex habitats). Permutation tests revealed that this effect was significantly different from random expectation (p= 0.03), where frogs with lower alpha diversity exhibit greater disturbance in bacterial community structure following exposure to *Ranavirus* than frogs with higher alpha diversity.

**Figure 4.**
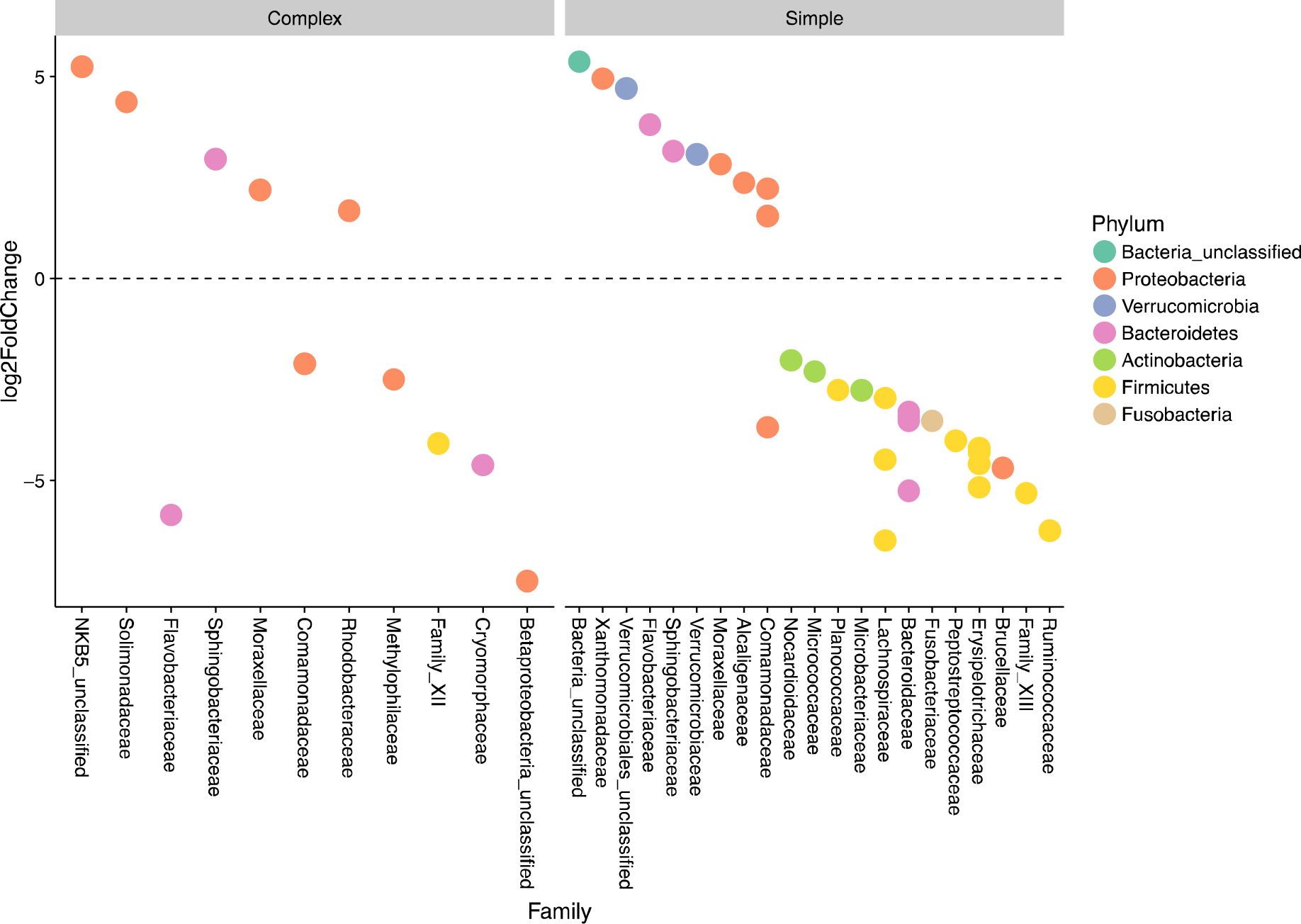
Change in relative abundance of OTUs in the common frog skin microbiome 48 hours post-exposure to ranavirus for individuals inhabiting complex (left panel) and simple (right panel) habitats. OTUs are coloured by by Phylum. Comparisons are to the sham-exposed individuals within the same habitat types. Exposure of individuals from simple habitats to *Ranavirus* resulted in a significantly greater number of changes in abundance compared to complex habitats, and involved more families and more phyla than for individuals from complex habitats.

### Links between habitat, microbiome diversity and survival following infection

Individuals in simple habitats exposed to *Ranavirus* exhibited higher rates of mortality (68.4%) than individuals in complex habitats exposed to ranavirus (52.2%; Fig. 5). Conversely individuals in both simple and complex habitats receiving a sham exposure showed limited mortality. When truncating the survival data to the day of exposure (n=88 individuals), the best-supported model contained effects of both habitat complexity and disease treatment on survival (Table 1). A model containing only disease treatment received marginally less support (ΔAICc = 0.22). Though the model containing the interaction between habitat and treatment was in the Δ6 AIC model set, it was a more complex version of a simpler model with better AIC support and so was removed under the nesting rule (Richards 2008). There was no difference between habitats in likelihood of exhibiting gross signs of disease (Binomial GLMM, mean probability of exhibiting signs of disease [95% credible intervals]: complex 0.48 [0.11,0.86]; simple 0.49 [0.1,0.89]; p_MCMC_= 0.92) or in severity of visible signs of disease (Ordinal GLMM, mean probability of being scored category 0 [95% credible intervals]; complex 0.49 [0.11,0.88]; simple 0.53 [0.11,0.93]; P_MCMC_= 0.87).

**Table 1.**
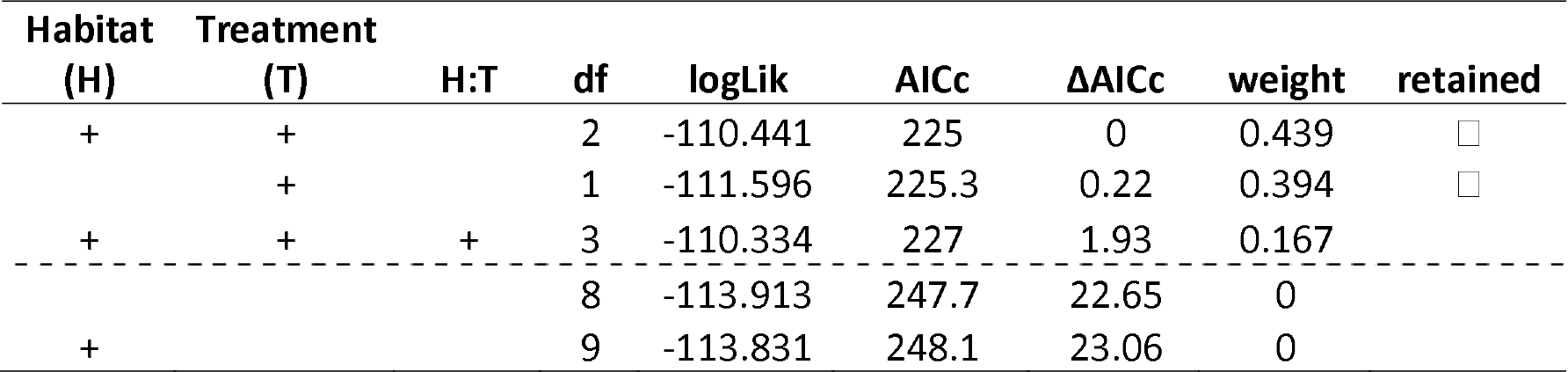
Model selection results from analysis examining the effects of habitat complexity (Habitat), disease treatment (Treatment) and their interaction (H:T) on the survival of common frogs. The best-supported model contained an effect of both Habitat and Treatment; individuals in simple habitats treated with ranavirus exhibited higher mortality rates than individuals in ranavirus-treated complex habitats, whilst there was limited mortality in both control groups. The second best-AIC model contained an effect of only ranavirus exposure (ΔAICc = 0.22). Although a model containing the H:T interaction was also in the Δ6 AICc model set, it is a more complex version of a model with better AIC support and so is removed under the nesting rule. Retained models are indicated with a tick mark in the ‘retained’ column. The Δ6AICc cutoff is indicated with a dashed line.

**Figure 5.**
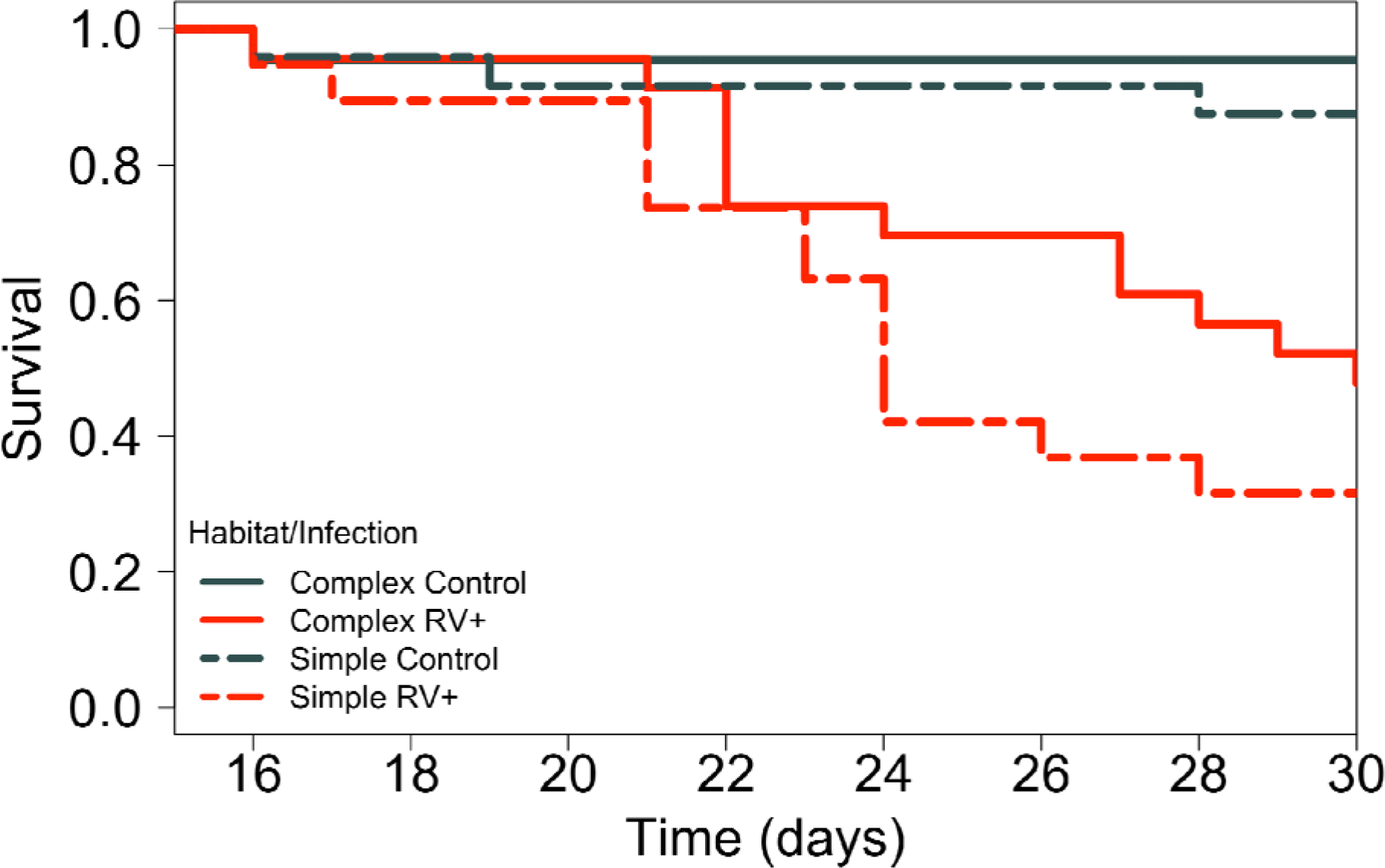
Survival data for 88 common frogs assigned to either simple (dashed lines) or complex (solid lines) habitat types, and infected with either ranavirus (RV+, red lines) or a sham control (grey lines). The best-supported model contained an effect of both infection and habitat, suggesting that Individuals in simple habitats infected with ranavirus showed higher mortality than ranavirus-infected individuals in complex habitats. All survival models controlled for block effects.

## Discussion

### Habitat complexity Predicts Host Microbiome complexity

Our results demonstrated that the structure and diversity of the amphibian skin microbiome is a function of the bacterial diversity present in the environment. These data support previous work implicating the role of the environment as a standing reservoir of microbes for host colonization and therefore driving host microbiome diversity in a range of taxa, including humans (Spor et al 2011; Cho & Blaser 2012; Lax et al 2014), reptiles (Hyde et al 2016; Kohl et al 2016), amphibians (Kueneman et al 2016; Longo & Zamudio 2017; Loudon et al 2014b; 2016) and corals (Pantos et al 2015). We detected many OTUs on frogs that were not detectable in the environment, especially for frogs inhabiting complex habitats, suggesting they may be present at extremely low abundance in the environment and difficult to detect. That these environmentally rare OTUs are enriched on amphibian hosts is suggestive of selection by the host of rare OTUs (Loudon et al 2016). These data comprise further evidence that host microbiome structure is not simply a direct function of random colonization from the environment (Walke et al 2014; Loudon et al 2016). Importantly, our data suggest that the extent of the enrichment of OTUs on amphibian skin relative to the environment is constrained by environmental complexity. Marked reductions in the reservoir of environmental bacteria may compromise the ability of hosts to enrich their skin microbiome with beneficial bacteria, which in turn may have consequences for host health (Kueneman et al 2016).

### Exposure to a Pathogen Disrupts the Host Skin Microbiome

Exposure to ranavirus elicited subtle but significant changes to the structure of the amphibian skin microbiome after 48 hours. Most notably, the magnitude of the effect was dependent on habitat complexity and indicated that low diversity host-associated bacterial communities are more susceptible to perturbation. Microbiome structure of infected individuals is a function of both initial microbiome state and perturbation caused by invasion of the pathogen (Jani & Briggs 2014; Walke et al 2015; Longo & Zamudio 2017), and these two contrasting effects must be accounted for when trying to understand how host microbiome structure relates to susceptibility to disease. That is, one cannot use microbiome composition of infected individuals to investigate links with disease susceptibility without knowing the relative contribution of the host/pathogen interaction to that observed state. Our results show that the magnitude of perturbation caused by exposure to a pathogen will not be equal for all hosts, and is instead a function of the initial state, measured here as diversity.

Measures of the magnitude of change in microbiome structure caused by disease are critical for understanding how invasion by a primary pathogen may facilitate secondary invasions. That the patterns of pathogen-mediated disruption to the microbiome differed between simple and complex habitats, indicates that more complex microbial communities are more resistant to perturbation than simpler communities. This concept aligns well with classical ecological theory (Shea & Chesson 2002; Costello et al 2012). So called ‘niche opportunity’ may be far greater in hosts harbouring species-poor bacterial communities and may result in far greater probability of successful invasions by opportunistic organisms (Shea & Chesson 2002). Conversely, destabilisation of community structure by an invading pathogen may release competitive pressure on bacterial species already present at low abundance and result in marked increases in relative abundance. The innate immune response to ranavirus in *Xenopus* can be rapid, with peak mononucleated macrophage-like cell activation occurring one day post infection (dpi) and natural killer (NK) cells peaking at three dpi (Morales et al 2010). This contrasts with the acquired immune response, where the T cell response peaks at six dpi (Morales et al 2010). We measured host microbiome change two dpi, suggesting that our results reflect the interaction between host innate immune response and microbiome diversity. Further work is required to assess whether the strength of the host innate immune response is modulated by skin microbiome diversity, but our data suggest that lower diversity communities are far more susceptible to perturbation by the immunological response that arises from pathogen exposure. Disruption of the host microbiome by pathogens of wild vertebrates is likely to be far more common than the existing literature suggests. The scarcity of studies directed at quantifying microbiome disruption by pathogens means we currently lack the ability to compare the magnitude of the perturbation effect among host species and both host and pathogen taxonomic groups.

### Covariance Between Environmental Heterogeneity and Survival

Data from the controlled infection experiment indicated that mortality in simple habitats was greater following exposure to *Ranavirus* than in complex habitats. Given the causal relationship between inhabiting complex habitats and significantly increased skin microbiome diversity, these data indicate a positive correlation between host microbiome structure and survival following exposure to a lethal pathogen. Studies have demonstrated that host-associated microbes can confer significant benefits to the host in the form of protection from infection and disease (reviewed in Ford & King 2016), and several studies have provided evidence consistent with a correlation between overall microbiome diversity and susceptibility to infectious disease and costs associated with host responses to pathogen exposure (Cariveau et al 2014; Kueneman et al 2016). For example, disruption of the normal microbiome by administration of antibiotics to laboratory mice can permit successful infection of *Clostridium difficile* (Theriot et al 2014), and loss of microbiome diversity in amphibians can increase susceptibility to the fungal pathogen *Bd* (Kueneman et al 2016). Notably, augmentation of low diversity skin microbiomes with key taxa from the more diverse wild-type microbiome can reverse the observed increase in susceptibility to a lethal amphibian pathogen like *Bd* (Kueneman et al 2016).

Variation in resistance to the impacts of infectious agents as a function of microbiome diversity could occur by at least three mechanisms. First, greater diversity of the microbiome may increase the chance of a bacterium with host-protective effects being present on the host, or able to persist on the host because of facilitation by other bacteria. Second, host protection may arise because community members act synergistically to provide immunity (Loudon et al 2014a). Third, microbiome complexity may indirectly modulate host traits linked to immunity, such as enhancing the production of anti-microbial peptides (AMPs) by the host (Rollins-Smith 2009). This is particularly relevant given previous work demonstrating that amphibian AMPs can inactivate ranavirus virions (Chincharr et al 2004) and most recently have been shown to be virucidal against the influenza virus (Holthausen et al 2017). Surprisingly, though survival was greater in complex habitats, this was not accompanied by a lower probability of exhibiting signs of disease. Collectively these data indicate that likelihood of infection was similar for both groups, but that differences in survivorship arose because host costs associated with either exposure to or infection with *Ranavirus* were less likely to reach lethal thresholds in animals with greater microbiome diversity. That we didn’t detect an effect of habitat on severity of signs of infection may mean the relationship between viral loads and signs of disease is non-linear, or may reflect low variance in disease state outcome among individuals, as very few individuals from either habitat type were scored the most severe disease category.

Our results have important implications for our understanding of the factors driving spatial and temporal heterogeneity in susceptibility to disease. Given the strong environmental component of determination of the amphibian skin microbiome (Longo & Zamudio 2017; this study), spatial or temporal variation in environmental complexity may also be mirrored in the complexity of the structure of the host microbiome, which in turn may drive variation in resistance to infection (Lam et al 2010; Chang et al 2016). Any activity that causes changes to the reservoir of environmental bacteria available to colonise hosts may in turn impact population-level susceptibility to disease (Longo & Zamudio 2017). Human activity can both increase the rate of pathogen spread (Price et al 2016) and alter the bacterial community dynamics of the environment (Sheik et al 2011; Deng et al 2012; Rodrigues et al 2013). Use of pesticides has been associated with increased prevalence of *Ranavirus* (North et al 2015), and could theoretically be mediated by disruption of the both environmental and host-associated bacterial communities. Our results highlight that the dramatic shifts in the structure and diversity of the environmental microbiome can be expected to have significant knock-on effects on the structure and diversity of the host-associated microbiomes of animals inhabiting those areas. The principal gaps in our understanding that remain are twofold; first, whether anthropogenically-mediated shifts in environmental and host microbiome structure can also modulate the strength of interaction between hosts and parasites, as demonstrated in this study and second, whether we can expect the degree of modulation to be uniform across the broad suite of pathogen life histories and modes of infection we observe in the wild.

## Acknowlegdements

XAH thanks Matt Gray for helpful discussion about the manuscript.

